# A mRNA Vaccine Encoding for a 60-mer G Glycoprotein Nanoparticle Elicits a Robust Neutralizing Antibodies Response Against the Nipah Virus

**DOI:** 10.1101/2024.08.05.606671

**Authors:** Pascal Brandys, César G. Albariño, Shilpi Jain, Irena Merenkova, Nicholas J. Shork, Aihua Deng, Martine Valière, Jens Herold

## Abstract

The Nipah virus is a zoonotic pathogen causing encephalitis and acute respiratory illness in humans with very high fatality rates. Here we report a novel messenger RNA vaccine that directly encodes for a nanoparticle displaying 60 head domains of the Nipah virus G (NiV G) glycoprotein that acts as a highly effective antigen. A vaccine encoding for the NiV G nanoparticle elicits high antibody titers against NiV G and a robust neutralizing antibody response with a pseudotyped Nipah virus system. We ultimately find that the proposed mRNA NiV G nanoparticle (mRNA NiV G-NP) provides a flexible platform for the development of vaccines that will likely be of great value in combatting future Nipah virus outbreaks.

## INTRODUCTION

The Nipah virus (NiV) is a paramyxovirus of the *Henipavirus* (HNV) genus discovered in 1999 and a bat-borne zoonotic pathogen causing encephalitis and acute respiratory distress syndrome in humans with a very high fatality rate. It was first isolated in pig farms around the Nipah river near Port Dickson, Malaysia ^1^. Natural hosts of the Nipah virus are large fruit bats of the *Pteropus* genus, which are geographically distributed across the Indian Ocean, South Asia, South-East Asia, Australia, and the Pacific Islands, including some of the most densely populated regions on Earth ^2^. Over the past two decades NiV has spilled over into humans almost annually in Bangladesh ^3, 4^ and India ^5^ (Bangladesh genotype) and has also caused outbreaks in Malaysia, the Philippines and Singapore ^6^ (Malaysian genotype). Of 248 Nipah virus cases identified in Bangladesh over 14 years, 82 were caused by person-to-person transmission corresponding to a reproduction rate of 0.33 ^4^. In August 2023 the Indian state of Kerala experienced its fourth outbreak in five years with reported human-to-human transmission and a fatality rate of 33% ^7^. To date, no approved vaccine or therapeutics for use in humans exist against Nipah virus infections. With the serious potential for larger epidemics in the future novel vaccine strategies against the Nipah virus are clearly required.

### Nipah Virus Neutralizing Antibodies Target the Nipah Virus Attachment G Glycoprotein

NiV is a single stranded, non-segmented, negative sense RNA virus with six structural proteins, including the attachment G and the fusion F envelope glycoproteins. NiV G is also known as the receptor binding protein (RBP). NiV entry into host cells requires fusion of the viral and host cell membranes through the concerted action of the G and F glycoproteins. NiV G is an oligomeric Type II transmembrane glycoprotein composed of a brief N-terminal cytoplasmic tail, a single transmembrane domain, and a sizable ectodomain with a stalk region and a C-terminal globular head domain ^8^. The NiV entry receptors at the surface of host cells are the transmembrane protein tyrosine kinases ephrin-B2 ^9^ or ephrin-B3 ^10^, despite these two receptor proteins only sharing 40% amino-acid identity. NiV G is a target of the humoral immune response ^11^ and serum neutralizing antibodies are correlated with protection in animals experimentally infected with NiV ^12^. In the African green monkey NiV causes a lethal respiratory and neurological disease that essentially mirrors the severe human disease. Vaccination of these primates with a recombinant subunit vaccine based on NiV G completely protects against subsequent NiV infection^12^. A single dose of a simian adenovirus-based vaccine encoding NiV G protects and prevents disease against a lethal challenge with NiV in Syrian hamsters ^13^. A vaccine based on a recombinant rabies vector expressing NiV G induces NiV G-specific neutralizing antibodies in mice ^14^. Adeno-associated virus vaccines expressing NiV G induce a potent and long-lasting antibody response in mice and protect against a lethal challenge in hamsters ^15^. A recombinant vesicular stomatitis vaccine expressing NiV G administered 7 days prior to exposure protects African green monkeys from lethal disease with survivors developing NiV G-specific neutralizing antibodies ^16^. A nucleoside-modified mRNA vaccine encoding soluble NiV G formulated into lipid nanoparticles induces NiV G-specific antibody response in pigs with high neutralization titers against NiV ^17^. The NiV G-specific m102.4 monoclonal antibody has been administered on an emergency compassionate use basis to 15 individuals with high-risk exposure to NiV infection and has completed a phase 1 clinical trial in Australia ^18^. The human neutralizing antibody NiV41-6 targeting a NiV G highly conserved epitope blocks the receptor binding interface and shows protective efficacy against a lethal NiV challenge in hamsters ^19^. All these results suggest that the NiV G glycoprotein is a prime immunogen candidate for rational antigen design of NiV vaccines.

## RESULTS

### Design of NiV G Antigen

NiV G crystal structure was determined in 2008 ^8^ and more recently the three-dimensional (3D) organization of NiV G protein was determined by cryo-electron microscopy (cryo-EM) in a model comprising the NiV G ectodomain from residues 96 to 602 ^20^. The NiV G protein forms a 200-Å-long and 120-Å-wide intertwined homotetramer. At the core of the 3D structure is an N-terminal stalk (residues 96-147) followed by a neck domain (residues 148-165) that connects with a linker region (residues 166-177) to a C-terminal head domain on each protomer (residues 178-602). The NiV G architecture adopts a distinctive two-heads-up and two-heads-down conformation. The NiV G head domain is the main target of antibodies neutralizing activity in sera elicited by vaccines using the full NiV G ectodomain tetramer for immunization ^20^ and the cell binding domain in NiV G was identified as residues 498-602 using a phage display system ^21^. The NiV G head domain is a suitable antigen for a nanoparticle vaccine as it comprises epitopes of the cell binding domain and retains its 3D structure when connected with a suitable linker region to an N-terminal multimerization domain. A NiV G head domain antigen based on the Malaysia or ancestral strain (GenBank: AF212302.2) is advantageous for the use of currently available antibody assays or neutralization assay systems using pseudotyped Nipah virus based on the same strain.

### Design of Glycosylated NiV G-Nanoparticle

High density display of glycosylated antigens onto protein nanoparticles enhances humoral immunity with improved access to lymph nodes, optimal packing and presentation of antigens ^22^. As demonstrated previously with various viral antigens ^23,24^ the self-assembling lumazine synthase (LS) from the hyper-thermophile *Aquifex aeolicus* is a suitable nanoparticle (NP) scaffold where 60 identical copies of the glycosylated antigen can be displayed with strict icosahedral 532 symmetry ^25^. Structural modeling suggested that with a suitable linker length the LS scaffold can sterically accommodate the glycosylated NiV G head domain, including four N-linked glycans (asparagine residues 306, 378, 417 and 529) that are highly conserved (Fig. 1). Fusing the NiV G head domain (residues 178-602) to LS results in a predicted spherical NiV G-NP 60-mer of a diameter of 40 nanometers mimicking the size, shape, multivalency and symmetry of many viruses to enhance immune response.

**Fig. 1.**
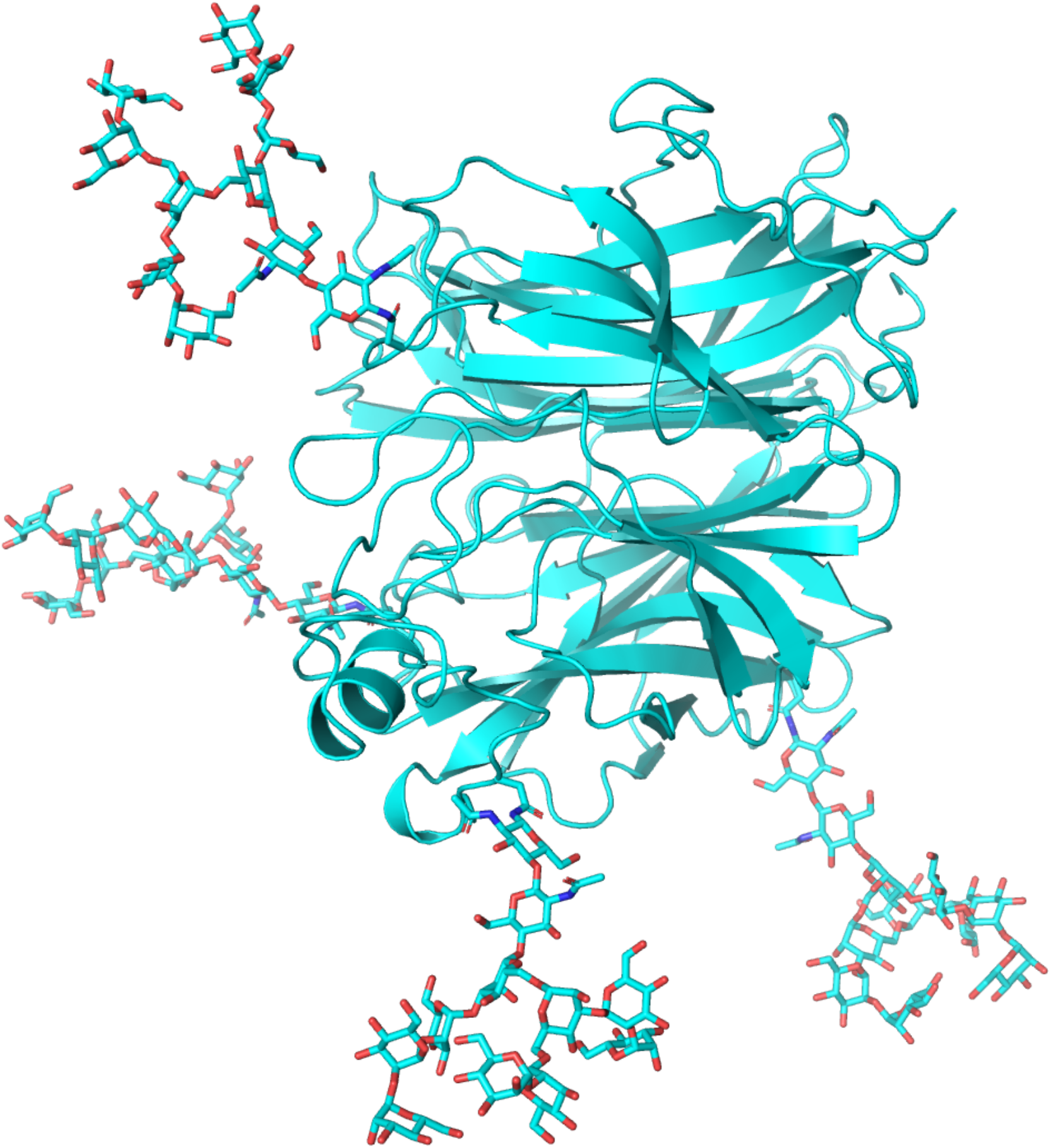
Cartoon representation of glycosylated NiV G head domain. (residues 178-602) with four N-linked glycans as predicted by AlphaFold2 (AF2).

### Design of mRNA Encoding for Glycosylated Nipah Virus G-Nanoparticle

We previously reported the design of a mRNA encoding for a 60-mer nanoparticle displaying the receptor binding domain of the severe acute respiratory syndrome coronavirus 2 (SARS-CoV-2) ^24^. The same design was used for the untranslated regions (UTR) and the translated LS region of a mRNA encoding for the NiV G head domain fused with LS, with a minimal 5’ UTR, a human beta-globin 3’UTR and a poly(A) tail of 80 nucleotides. The mRNA was not modified, neither with guanosine/cytidine (G/C) content modification nor with chemical modification.

### Design of Vectors for In Vitro Transcription

As described previously a site for linearization with BsmBI was inserted 3’ to the poly(A) tail, resulting in a free 3’ end after the poly(A) tail ^24^. A construct with the minimal 5’ UTR, the human IL-2 signal sequence, a nucleotide sequence encoding the NiV G head domain fused with LS, the human beta-globin 3’UTR, a poly(A) tail of 80 adenosine residues and the BsmBI restriction site was cloned into a pUC19 vector. The supercoiled pUC19 DNA was upscaled, linearized with BsmBI and purified. In vitro transcription was performed with T7 RNA polymerase in a 5 ml reaction. The mRNA was capped with a cap 1 structure on the 5’ end by vaccinia 2’-O-methyltransferase enzymatic capping. The capped mRNA was purified by reverse phase chromatography followed by tangential flow filtration (TFF). Final yield of purified mRNA was 5.51 g/l of IVT reaction.

### Vaccine Formulation and Delivery

The mRNA-NP vaccine was formulated as described previously ^24^. The small arginine-rich DNA-binding nuclear protein protamine is widely used to deliver different types of RNA in vitro and in vivo. mRNA complexed with clinical grade polycationic protein protamine shows significantly improved cytosolic delivery capacity and results in a self-adjuvanted vaccine ^26^. The mRNA was complexed by addition of clinical grade protamine to the mRNA at a mass ratio of 1:5. The vaccine was prepared on each injection day with final total mRNA concentration of 840 μg/ml. As previously reported needle-free injection improves the efficacy of mRNA-NP vaccines ^24^ and the vaccine was injected into the caudal thigh muscle of mice with a needle-free injection system (Tropis modified for mouse injection, PharmaJet).

### Immunization with mRNA NiV G-NP Vaccine Elicits High Serum Anti-NiV G Antibody Titers in Mice

To investigate the prophylactic potential of the mRNA NiV G-NP vaccine against the Nipah virus we gave to one group (N=10) of mice a prime/boost regimen of the mRNA NiV G-NP vaccine. CB6F1/J female mice 7 weeks old were primed at week 0 and boosted at week 3 with the same dose of 42μg/50μl by intramuscular injection at the caudal thigh with a needle-free injection system. Blood was collected and serum prepared on weeks 0 (prior to prime), 3 (prior to boost) and 6.

To determine if the mRNA NiV G-NP vaccine elicited neutralizing antibodies directed to the NiV G head domain we analyzed samples of weeks 3 and 6 with the anti-NiV G antibody detection enzyme-linked immunosorbent assay (ELISA). The sandwich ELISA detects any antibodies targeting the Nipah virus G glycoprotein (Malaysia strain). A positive control mouse anti-NiV G monoclonal antibody indicated a sensitivity of the assay below 1 ng/ml of antibody and optical density (OD) a linear function of antibody concentration in the 1-100 ng/ml range (Fig. 2). In all 10 mice immunized with mRNA NiV G-NP vaccine a high concentration of antibodies (>100 ng/ml) targeting NiV G was detected in week 3 and week 6 sera, with OD = 4.0 in all week 3 and week 6 mouse sera samples at volume dilution ratios of 1:10, 1:100, 1:1000 and 1:5000, and in 10/10 week 6 and 9/10 week 3 mouse sera at a volume dilution ratio of 1:10000. At a volume dilution ratio of 1:50000 OD results were significantly different between week 3 (mean 1.39, sd 0.46) and week 6 (mean 3.79, sd 0.33) mouse sera (Fig. 2 B) and, as expected, near the negative control value in all week 0 mouse sera prior to immunization (mean 0.33, sd 0.13). At a volume dilution ratio of 1:100000 all week 6 sera were in the linear part of the positive control curve with OD ranging from 1.33 to 3.57 (mean 2.54, sd 0.72) (Fig. 2 C) and an extrapolated average concentration of anti-NiV G antibodies >1 mg/ml in mouse sera. Between week 3 and week 6 the concentration of anti-NiV G antibodies in mouse sera increased on average 12-fold (Fig. 2 D).

**Fig. 2.**
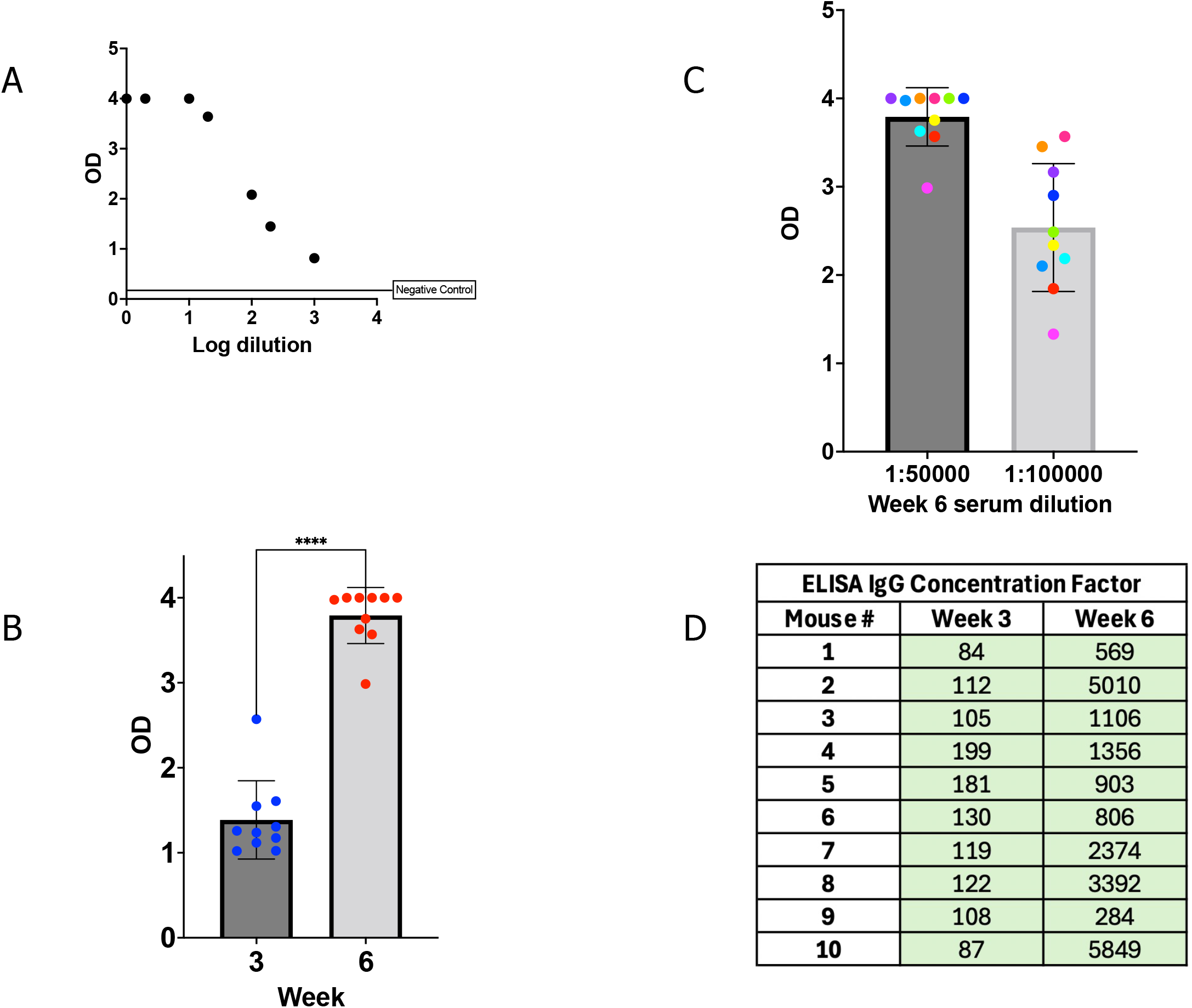
ELISA. **A) Positive control:** Optical density (OD) values of anti-NiV G antibody at 7 serial volume dilutions starting with a concentration of 1 *µ*g/ml. **B) OD of week 3 and 6** mouse serum samples at 1:50000 volume dilution (mean values of duplicate wells). **C) OD of week 6** mouse serum samples at 1:50000 and 1:100000 volume dilutions (mean values of duplicate wells). **anti-NiV G IgG ELISA concentrations** in the samples tested at 1:100000 volume dilution. The concentration factor is the concentration of the positive control (*µ*g/ml) equal to the concentration of anti-NiV G IgG in the sample as measured by OD.

### Immunization with mRNA NiV G-NP Vaccine Elicits High Serum NiV Neutralizing Titers in Mice

To determine if the mRNA NiV G-NP vaccine elicited NiV neutralizing antibodies we analyzed immunized mouse sera samples of weeks 3 and 6 with a novel neutralization assay developed for use in a biosafety level 2 laboratory ^27^. The assay uses a recombinant vesicular stomatitis virus expressing NiV G and the fluorescent reporter protein ZsG (rVSV-ΔG-NiV). The recombinant virus propagates as a replication-competent virus in a cell line constitutively expressing the NiV F fusion protein. The neutralization activity observed in the assay using the pseudotyped virus rVSV-ΔG-NiV G is comparable with the neutralization activity observed using a fully infective NiV in a biosafety level 4 laboratory. In all 10 mice immunized with mRNA NiV G-NP week 3 and week 6 sera neutralized rVSV-ΔG-NiV G in a dose-dependent manner. All 10/10 week 6 sera samples and 7/10 week 3 sera samples neutralized the pseudotyped virus in an efficient manner (Fig. 3). 50% effective inhibition concentration (IC_50_) values of mouse sera against rVSVΔG-NiV-G at week 6 ranged from 1120 to 5302 (mean 1984, sd 1310) (Fig. 3 B). For reference IC_50_ values with the same assay were previously reported^27^, testing 14 positive human samples collected during NiV outbreaks in Bangladesh (mean 681, sd 550). For comparison with another NiV G vaccine in mice, 45 days post-vaccination with a recombinant rabies virus vector expressing NiV G, average fluorescence reduction neutralization assay 50 (FRNA_50_) were below 200 in one group vaccinated with one dose of live viral particles (N=10) and below 400 in another group vaccinated with prime-boost of chemically inactivated viral particles (N=10)^14^. For comparison with other NiV G vaccines in various animal models, 35 days post-vaccination with a recombinant vesicular stomatitis vaccine expressing NiV G average 50% plaque reduction neutralization tests (PRNT_50_) were below 256 and below 1024 for two groups (N=5) of African green monkeys that were protected from the lethal disease^16^, 39 days post-vaccination with a replication-deficient simian adenovirus vector encoding NiV G 50% inhibition values in virus neutralization assays were below 80 for Syrian hamsters receiving a single dose (N=9) and below 240 for hamsters receiving prime-boost (N=10), all these animals survived a lethal NiV challenge with no sign of the disease^13^, 42 days post prime-boost vaccination with a mRNA vaccine encoding NiV G average NiV neutralization titers were below 1000 in a group of pigs (N=6)^17^. As expected, no neutralization activity, even at lower dilutions, was observed in mouse sera samples collected at week 0 prior to immunization (Fig. 3). For 7/10 mouse sera at week 3 50% effective inhibition concentration (IC_50_) ranged from 117 to 546 (mean 249, sd 144) and concentration-response curves in Vero-Fco cells were comparable to the positive control serum curve (Fig. 4), indicating efficient neutralization activity 3 weeks after the first dose and prior to the second dose. On average neutralization titers increased 11-fold between week 3 and week 6, and the average ELISA concentration of anti-NiV G antibodies increased 12-fold.

**Fig. 3.**
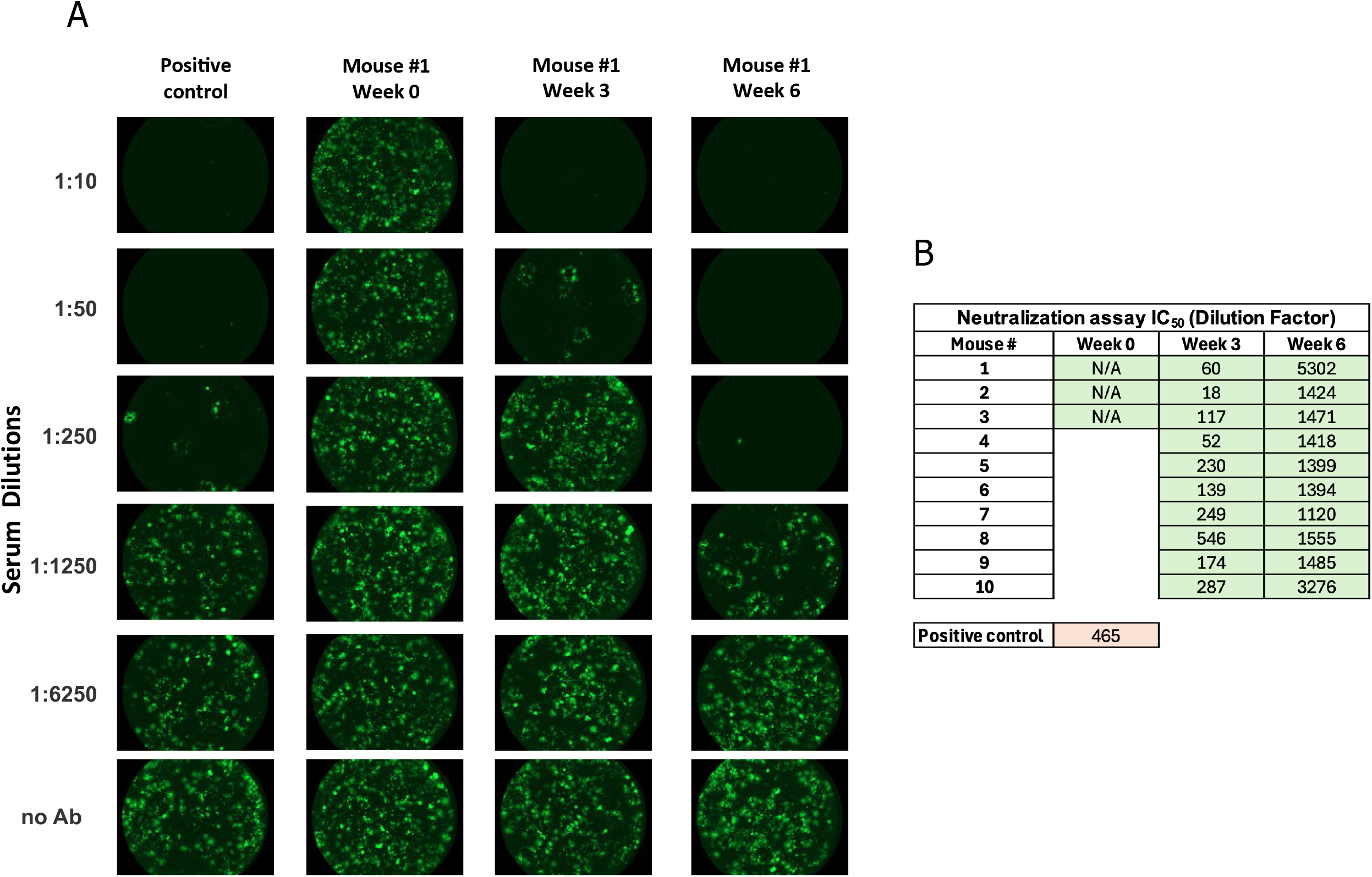
Neutralization assays. **A) Neutralization assays using rVSVΔG-NiV-G:** Vero-Fco cells were treated with serial 5-fold dilutions of indicated mouse sera mixed with 100 TCID_50_ of the recombinant virus (rVSVΔG-NiV-G). After 72 h, fluorescence signal was measured using Synergy. Representative images of the neutralization activity of anti-NiV G glycoprotein in cells infected with recombinant virus. **B) 50% effective inhibition concentration (IC_50_) values** of each indicated mouse serum against rVSVΔG-NiV-G. The positive control is serum of a hamster that survived a Nipah virus challenge.

**Fig. 4.**
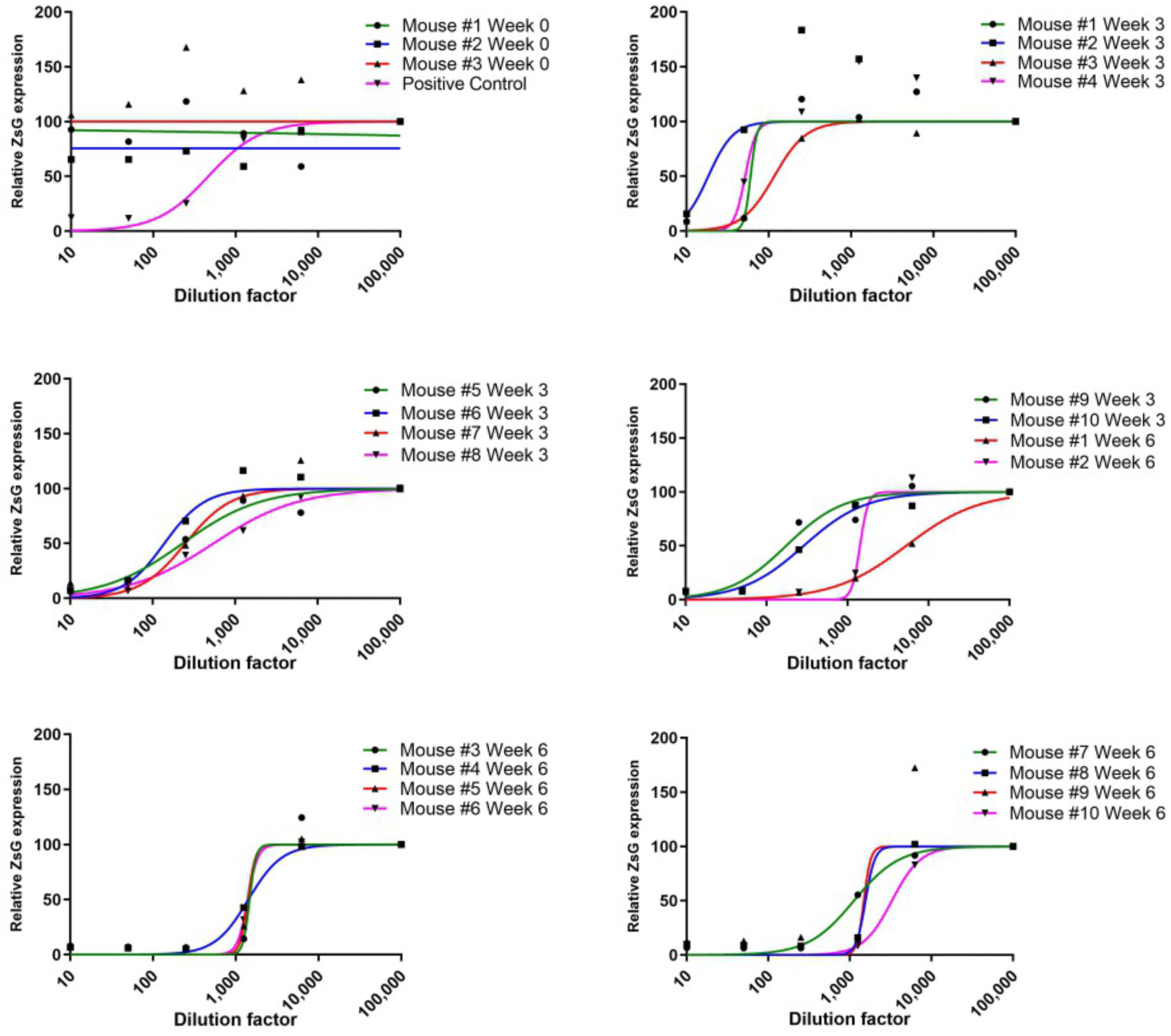
Concentration-response curves in Vero-Fco cells: Vero-Fco cells were infected with recombinant virus after neutralization with mouse sera. Relative fluorescence units (RFU) are shown on the y-axis. Each point on the graph represents mean values, and error bars indicate standard deviations of triplicate wells.

## DISCUSSION

In this study we demonstrated that the mRNA NiV G-NP vaccine elicits a robust anti-NiV G humoral response against the G glycoprotein head domain with high serum antibody titers. We also demonstrated that the vaccine elicits a robust NiV neutralizing antibody response with high serum NiV neutralizing titers using a pseudotyped Nipah virus system. All these results were obtained with assays based on the Malaysia strain (GenBank: AF212302). The Bangladesh strain (GenBank: AY988601) differs from the Malaysia strain by 14 mutations in the 425 residues of the NiV G head domain (Table 1). Further cross-neutralization assays of the Bangladesh strain will assess the effect of this limited genetic drift on NiV neutralization with sera elicited by the mRNA NiV G-NP vaccine. In this study we also demonstrated comparable neutralization activity between 7/10 immunized mouse sera at week 3 and serum of a hamster surviving NiV infection. This is indicative of potential efficacy of the mRNA NiV G-NP vaccine 3 weeks after the first dose, opening the perspective of emergency use of the mRNA NiV G-NP vaccine for immunization during Nipah virus outbreaks.

**Table 1:**
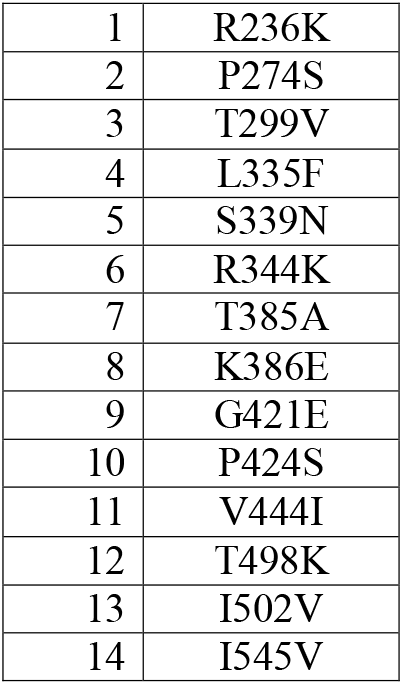
The Bangladesh strain (GenBank: AY988601) differs from the Malaysia strain (GenBank: AF212302) by 14 mutations in the 425 residues (178-602) of the NiV G head domain.

|  |  |
| --- | --- |
| 1 | R236K |
| 2 | P274S |
| 3 | T299V |
| 4 | L335F |
| 5 | S339N |
| 6 | R344K |
| 7 | T385A |
| 8 | K386E |
| 9 | G421E |
| 10 | P424S |
| 11 | V444I |
| 12 | T498K |
| 13 | I502V |
| 14 | I545V |

### mRNA NiV G-NP Vaccine Can Withstand the Evolution of the Nipah Virus

The evolutionary rate of NiV was estimated at 4.64 × 10^−4^ nucleotide substitutions/site/year for the Bangladesh genotype ^28^ and is relatively slow as compared with other RNA viruses, e.g. 1.92 × 10^−3^ substitutions/site/year for human respiratory syncytial virus B ^29^ and 7.30 × 10^−4^ substitutions/site/year for SARS-CoV-2 before the emergence of variants of concern ^30^. This nucleotide substitution rate would translate into less than one amino-acid substitution in the 425 residues of the NiV G head domain every 5 years. Alternatively, the Bangladesh and Malaysian genotypes diverged about 75 years ago ^28^ with 14 accumulated amino-acid substitutions in the NiV G head domain, or on average one amino-acid substitution every 5.3 years, a nearly identical estimate. Interestingly, few of the NiV G mutations in known NiV strains are antibody escape mutations. A recent study using a pseudotyped virus deep mutational screening library of NiV G to map NiV escape mutations for six neutralizing antibodies that target a diversity of epitopes on the NiV G head domain found only one amino-acid mutation in natural Nipah strains (P274S in Bangladesh genotype) ^31^. Assuming limited human-to-human transmission and antigenic escape of NiV, and the current evolutionary rate of the NiV G head domain, the mRNA NiV G-NP vaccine could maintain its efficacy over a decade or more. In the long term and if necessary for future outbreaks one major advantage of the mRNA NiV G-NP platform will be to allow the rapid development and manufacturing of updated vaccines.

### Understanding and Addressing the Diversity of the Nipah Virus with Optimized mRNA NiV G-NP Vaccine

In NiV-affected regions of the state of Kerala an overall seroprevalence of 21% was observed in 272 *Pteroptus medius* bats tested by anti-Nipah bat immunoglobulin G enzyme-linked immunosorbent assay (ELISA) during 2023. The partial N gene retrieved showed more than 99% similarity with earlier reported NiV genomes from Kerala belonging to the cluster of NiV Indian sequences ^32^. The sequences of the NiV strain from the latest 2023 Kerala outbreak also clustered into the same Indian clade ^7^. A mRNA NiV G-NP vaccine based on the NiV G head domain sequence of this Indian clade can be developed with the objective to prevent NiV spillover from bats to the human population in India. NiV does not exhibit significant genetic variation in Bangladesh ^28,33^ also facilitating the design of a mRNA NiV G-NP vaccine for this country. In a few sites in Bangladesh and Cambodia where genomic surveillance has been concentrated most clusters have been identified but only ∼15% of overall NiV diversity has been identified ^34^. Recent initiatives for the sequencing of additional NiV strains ^35^ should improve the understanding of the diversity of the Nipah virus in many endemic countries and facilitate the design of a universal monovalent or multivalent mRNA NiV G-NP vaccine candidate. NiV infections in humans will likely remain sporadic and unpredictable, preventing a traditional pathway of clinical development for the mRNA NiV-NP vaccine. After a phase 1 safety and immunogenicity study, additional clinical studies of the mRNA NiV G-NP vaccine candidate in exposed individuals on a compassionate basis should be pursued as a priority.

## MATERIALS AND METHODS

### Mice

CB6F1/J female mice were purchased from The Jackson Laboratory (SN 100007). During the immunization period, mice were kept in individually ventilated cages in the animal facility of BTS Research in San Diego, CA, USA. Mice were fed standard chow ad libitum and kept in a 14:10 light:dark cycle.

### mRNA Encoding NiV G-NP

mRNA encoding the NiV G head domain fused with LS was designed with a minimal 5’ untranslated region (UTR) of 14 nucleotides, a human beta-globin 3’UTR, 3’ polyadenylation with a poly(A) tail of 80 nucleotides, and unmodified nucleosides.

### In Vitro Transcription of mRNA

A construct with the minimal 5’ UTR, the human IL-2 signal sequence, a nucleotide sequence encoding NIV G head domain fused with LS, the human beta-globin 3’UTR, a poly(A) tail of 80 adenosine residues and the BsmBI restriction site was generated and cloned into a pUC19 vector by Officinae Bio in Venice, Italy. The supercoiled pUC19 DNA was upscaled, linearized with BsmBI, and purified. In vitro transcription was performed with T7 RNA polymerase in a 5 ml reaction. The mRNA was capped with a cap 1 structure on the 5’ end by vaccinia 2’-O-methyltransferase enzymatic capping. Capped mRNA was purified by reverse phase chromatography using POROS^™^ resin (ThermoFisher) followed by tangential flow filtration (TFF). Final yield of purified mRNA was 5.51 g/l of IVT reaction. RNA integrity was analyzed by denaturing PAGE. mRNA sample and ladder were denatured at 70°C for 2 minutes and kept on ice, then 200 ng of mRNA mixed with diluent marker was loaded on gel.

### Formulation

The mRNA was complexed by addition of clinical grade protamine to the mRNA at a mass ratio of 1:5. The vaccine was prepared on each injection day with final total mRNA concentration of 840 mg/l. mRNA integrity prior to and after formulation was analyzed on 1% agarose gel with SYBR safe stain (ThermoFisher).

### Anti-NiV G Antibody Detection ELISA

Sera samples of week 3 and 6 were analyzed with anti-NiV G antibody detection ELISA. The sandwich ELISA detects any antibodies directed to the G glycoprotein (Malaysian strain). The ELISA has four key components: a nickel chelate-coated 96-well plate (Pierce ThermoFisher, Cat #15442), a NiV G glycoprotein (residues 71-602, Malaysian strain accession# Q9IH62-1) produced in HEK293 cells with hexahistidine tag on the C-terminus used for plate capture (AcroBiosystems, Cat #GLN-N52H3-50), a horseradish peroxidase (HRP) conjugated goat anti-mouse immunoglobulin G fragment crystallizable region (IgG Fc) secondary antibody (Pierce ThermoFisher, Cat #SA510276), and a mouse monoclonal antibody specific for NiV G glycoprotein (JB3) (The Native Antigen, Cat #MAB12308-100) used as positive control.

First, the nickel chelate-coated plate was pre-washed. The plate was then coated with NiV-G protein by adding 0.5μg of polyhistidine-tagged NiV-G per well and incubating with shaking for 1 hour at room temperature (20-25°C). After washing steps, the positive control with a series of dilutions, the negative control (phosphate buffered saline, PBS) and the sera samples at serial dilution volumes 1:10 to 1:100000 with PBS were added in duplicate to the wells and incubated for 1 hour at room temperature. After washing steps antibodies in mouse sera targeting NiV G remained bound to the plate. The HRP-conjugated secondary anti-mouse IgG Fc antibody was added to detect the mouse antibodies on the plate and incubated for 1 hour at room temperature. After washing steps 3,3’,5,5’-tetramethylbenzidine (TMB) solution was added and the plate was incubated in dark at 20°C for 15 min. The reaction was quenched by adding 0.16M sulfuric acid and the final solution was read immediately at 450 nm in a microtiter plate reader. The absorbance of the sample was dependent on the concentration of anti-NiV G antibodies in the sample as indicated by the positive control curve (Fig. 2 A). The anti-NiV G IgG concentration factor in the sample, defined as equal to the concentration of the positive control (μg/ml) as measured by OD, was calculated by simple linear regression applied to the linear part of the positive control curve.

### Generation of Recombinant Pseudotyped Virus rVSV-ΔG-NiV G

Recombinant rVSV-ΔG-NiV-G, expressing NiV G protein and fluorescent reporter protein ZsG, was generated using NiV Malaysia strain (GenBank: AF212302) as described previously ^27^. Briefly, VSV glycoprotein G was replaced with NiV G and ZsG was inserted upstream of the VSV N (nucleoprotein) gene, separated from N by the self-cleaving 2A porcine teschovirus (P2A) sequence. To generate helper plasmids for rescue, VSV G (glycoprotein), VSV L (polymerase), VSV N, VSV P (phosphoprotein), and T7 RNA polymerase genes were cloned into a standard pol II expression vector (pCAGGS). To rescue rVSV-ΔG-NiV G, different concentrations of full-length pVSV-ΔG-NiV G, pCDNA-T7, pCDNA-VSV L, pCDNA-VSV N, pCDNA-VSV P, and pCDNA-VSV G were transfected into Huh-7 cells, and the supernatants were blind-passaged twice onto Vero-Fco cells, a cell line constitutively expressing NiV F. Recombinant viruses were titered using a standard 50% tissue culture infectious dose (TCID_50_) using the Reed and Muench method in Vero-Fco cells. Viral sequences were confirmed by next generation sequencing (NGS) on a MiniSeq platform (Illumina), and sequences were analyzed using CLC genomics workbench v 20.0 (Qiagen)(Genbank: PQ141688).

### Neutralization Assays

Neutralization assays were performed on week 0 mouse sera collected before immunization, week 3 mouse sera collected before the second dose, and week 6 fully immunized mouse sera. Vero-Fco cells have been described previously ^36^. Briefly, a codon-optimized sequence encoding the Fusion protein of NiV strain Malaysia (Genbank: AF212302) was inserted into the cell genome using the Flp-In system (Invitrogen). For the neutralization assay, 10,000 cells/well were seeded onto 96-well plates one day prior to the experiment. Serial 5-fold dilutions of mouse serum samples were mixed with 100 TCID_50_ of rVSV-ΔG-NiV G. After the virus/sample mixture was incubated for 1 h at 37°C, it was added to Vero-Fco monolayers in triplicate. Cells were incubated for 72 h at 37°C and the fluorescence signal was measured using a plate reader (BioTek). Data were analyzed using GraphPad Prism v9 (GraphPad Software). Fluorescence values of ZsG expression were normalized to fluorescence in non-infected cells. The resulting normalized values were used to fit a 4-parameter equation to semi-log plots of the concentration-response data. Dilution of the sera samples inhibiting 50% of ZsG expression (IC50) was calculated by interpolating the values.

### Immunization

One group (N=10) of immunized mice was primed at week 0 and boosted at week 3 with the same dose of 42μg/50μl by intramuscular injection at the caudal thigh under 1-5% isoflurane anesthesia with a needle-free injection system (Tropis® injector prototype modified for mouse injection, PharmaJet). Blood was collected by retro-orbital sinus puncture at week 0 (prior to prime) and 3 (prior to boost), and terminal cardiac puncture at week 6, and serum samples were prepared.

### Computational Modeling and Artificial Intelligence

Structural models of NiV G head domain were generated with AlphaFold2 (AF2) version 2.3.2 (Google DeepMind) protein folding deep learning system.

### Statistical Analysis

Statistical analyses were performed using Prism v9.3.1 (GraphPad Software). Statistical differences between groups in datasets with one categorical variable were evaluated by two sample t-test (2 groups) or one-way ANOVA (more than 2 groups) corrected for multiple comparisons unless indicated otherwise. Correlations between two variables were evaluated with simple linear regression and Pearson correlation coefficient unless specified otherwise. p values ≤ 0.05 were considered significant with the following reporting style in the figures:

ns p>0.05

* p ≤ 0.05

** p ≤ 0.01

*** p ≤ 0.001

**** p ≤ 0.0001

## DATA AVAILABILITY STATEMENT

The original contributions presented in the study are included in the article and supplementary material. Further inquiries can be directed to the corresponding author.

## ETHICS STATEMENT

Animal welfare for the mouse studies in the U.S. was conducted in compliance with the U.S. Department of Agriculture (USDA) Animal Welfare Act (9 CFR Parts 1, 2, and 3) and the Guide for the Care and Use of Laboratory Animals, 8th Edition, National Academy Press, 2011, Washington, DC, USA. The studies design and animal usage were reviewed and approved by the Institutional Animal Care and Use Committee under number 23-031 for compliance with regulations prior to study initiation.

## AUTHOR CONTRIBUTIONS

PB conceived, designed, supervised and analyzed the experiments and wrote the paper; CA designed and performed the neutralization assays; SJ provided the stock of recombinant VSV and cells for its propagation; IM designed and performed the vaccine formulation and prepared the ELISA; NS performed statistical analysis and edited the manuscript; AD performed and analyzed the ELISA; MV provided formulation material; JH assisted with study design and supervision and reviewed the manuscript. All authors contributed to the article and approved the submitted version.

## ACKNOWLEDGMENTS

The findings and conclusions in this report are those of the authors and do not necessarily represent the official position of the Centers for Disease Control and Prevention.

## CONFLICT OF INTEREST

PB and JH are the founders and owners of Phylex BioSciences, Inc., the company that holds IP related to mRNA NiV G-NP vaccine; PB is a named inventor on NiV G-NP vaccine patent; AD is employed by BTS Research. The remaining authors declare that the research was conducted in the absence of any commercial or financial relationships that could be construed as a potential conflict of interest.

